# Association of SYNE2 variants in accelerating the progress of DYT1 early-onset isolated dystonia

**DOI:** 10.1101/807891

**Authors:** Feng-Chin Lee, Steve Wu, Chih-Sin Hsu, Shih-Ming Huang, Jau-Shyong Hong, Chih-Fen Hu

**Author notes:** Correspondence: Chih-Fen Hu.

## Abstract

DYT1 early-onset isolated dystonia (DYT1 dystonia), a rare autosomal dominant (AD) primary dystonia, is categorized as a monogenic disease. While it is a well-known AD inherited disease, the relatively low penetrance rate implicates potential modifiers in play for disease progression. In this report, an affected individual with *TOR1A* gene (c.907_909delGAG, p.E303del) variant, was identified along with three additional AD carriers in the family. Since we failed to find the second hit variant from TOR1A (D216H, F323_Y328del and F205I) and major binding proteins, including TOR1AIP1 and 2 or HSPA8 proteins, subsequent whole exome sequencing on the patient, the carriers and a non-carrier family member were performed to screen for candidate modifiers of TOR1A (E303del). The result reveals that this patient distinctly carries one copy of *TOR1A* gene (c.907_909delGAG, p.E303del) and one or two copy of *SYNE2* gene (c.1721T>C, c.12001T>C, and c.12002G>A), encoding I574T, W4001R, and W4001Ter variants. We propose that these SYNE2 variants are linked to earlier disease onset in this patient by impacting the protein-protein interaction between TOR1A and SYNE2. Our study suggests *SYNE2* gene maybe a culprit to lower the threshold for DYT1 dystonia progression and provides one novel gene target for further screening diagnosis of DYT1 dystonia.

## Introduction

DYT1 (dystonia 1 protein) early-onset isolated dystonia (DYT1 dystonia) is a rare hereditary form of dystonia. It follows a pattern of autosomal dominant (AD) inherited movement disorder without other neurological symptoms, signs, and secondary causes. It often begins in childhood and adolescence (< 28 years old) and the symptom commonly starts from lower limbs and frequently propagates to other body regions when the disease progresses (1). The disease frequency in the Ashkenazi Jewish population is calculated to be 1:3000–1:9000 while it is approximately five times lower in the non-Jewish population (2). There is no known frequency in Taiwan-based population so far, even not available in Asia area. DYT1 dystonia is a severe form of primary torsion dystonia with a penetrance rate around 30∼40% (3). This disease has a strong hereditary predisposition but lacks a distinct neuropathology. In isolated dystonia, cognition and intellectual abilities remain intact despite the presence of significant movement abnormalities (4).

Many patients diagnosed as DYT1 dystonia is originally caused by a mutation in the *TOR1A* gene that locates at chromosome 9q34.11 and encodes TOR1A, an ATP-binding protein. A three-base pair deletion (c.907_909delGAG, p.E303del, rs80358233) resulting in loss of a glutamic acid residue in the TOR1A protein is identified in most affected individuals (5). The TOR1A protein can be found in the endoplasmic reticulum and the nuclear envelope of most cells, including those of the central nervous system (CNS). The molecular and cellular processes in which TOR1A is involved include the interactions between cytoskeleton and membrane and the important functions of endoplasmic reticulum and nuclear envelope (6). However, the function of TOR1A and how *TOR1A* gene pathogenic variants lead to dystonia remains largely unknown. Moreover, the markedly decreased penetrance poses a momentous challenge for diagnostic testing and genetic counseling, but also provides strong viewpoint for the existence of additional genetic modifiers which influence penetrance and variability of the disease (7).

Here, we report a boy who first exhibited waddling gait followed by fast development of typical dystonia. We surveyed the patient’s candidate genetic alterations first and found a deletion of residue 303 glutamic acid in the TOR1A (E303del). Beyond GAG-deletion, we initially screened for protective TOR1A (D216H) and pathogenic TOR1A (F323_Y328del and F205I) (8,9) variants, but the results were all normal alleles in the case. Then, we targeted the variants from the major binding partners of TOR1A, including TOR1AIP1 and 2 or HSPA8 (10,11), but the variants from the *TOR1AIP1* gene have high allele frequency in the dataset which implies that they are less likely as modifiers in our case. Therefore, we performed whole exome sequencing (WES) among the patient, his parents, and two additional AD carriers to find the candidate variants. The result of WES analysis revealed three single nucleotide variants (SNV) from the *SYNE2* gene (c.1721T>C (p.I574T), c.12001T>C (p.W4001R), c.12002G>A (p.W4001Ter)) and we propose that these variants may link to earlier disease onset in this patient.

## Methods

### DNA Samples from the Patient and Family Members

Total 10 individuals, including 1 patient and 9 family members, were investigated in the study. After acquiring written informed consent from all individual participants included in the study, the genomic DNA were extracted from blood leukocytes using MagPurix^®^ automated DNA extraction system. Detailed clinical information was obtained from corresponding clinicians and medical records. The experimental protocols were approved by the Institutional Review Board of Tri-Service General Hospital, National Defense Medical Center (1-107-05-164).

### Sanger Sequencing

Two variants of *TOR1A* gene and one variant of *SYNE2* gene were tested by published primers for PCR amplification across the critical region of desirable exons territory. (1. *TOR1A* (c.646G>C), Forward: TAATTCAGGATCAGTTACAGTTGTG, Reverse: TGCAGGATTAGGAACCAGAT; 2. *TOR1A* (c.907_909delGAG), Forward: GTGTGGCATGGATAGGTGACCC, Reverse: GGGTGGAAGTGTGGAAGGAC; 3. *SYNE2* (c.1721T>C), Forward: CCTGGGAAAATTCTTGCTTTC, Reverse: ATGTGCGTGTTTGACCATGT).

### Whole Exome Sequencing

Purified genomic DNA was randomly fragmented to size between 150 and 200 bp using Covaries S220. SureSelectXT Human All Exon V6 was used to perform exome capture for further sequencing. Whole exome sequencing was performed using an Illumina HiSeq 6000 platform with 150-base paired-end reads and output data is output data is up to 10Gb per sample. Sequencing data were analyzed following GATK best practices workflows (https://software.broadinstitute.org/gatk) for germline SNV and indel calling (12). Briefly, using Burrows–Wheeler Aligner to perform alignment with human hg38 reference genome. After alignment, remove duplicate performed by picard software, and using GATK to perform local realignment and base quality recalibration. SNVs and indels was identified using GATK-HaplotypeCaller. GATK-SelectVariants function were used to generate subsets of variants for further analyses. These variants were validated by manually viewing in *Integrative Genomics Viewer* followed by annotation with database, include refGene, clinvar_20170905, avsnp150, dbnsfp33a, gnomad_genome, dbscsnv11 in ANNOVAR software. Final candidate variant was confirmed by Sanger sequencing (Genomics^®^, Taipei, Taiwan). Sequencing data has been deposited to the GenBank databases under SRA accession: PRJNA523662 (https://www.ncbi.nlm.nih.gov/sra/PRJNA523662).

## Results

### Clinical Observations

The patient, without any major disease history in the past, had an initial presentation of waddling gait at 7 years old and the symptom progressed to limbs tremor within a few months. He is the only one affected with dystonia in the family. The other family members are all free from dystonia related neurological disorders. At the early phase of disease, limbs tremor, pronation of the upper limbs and waddling gait were noticed in full consciousness while symptoms faded away during sleep. Cognition, communication and mental acuity were all preserved. Over time, head tilt, scoliosis, kyphosis, repetitive and active twisting of limbs appeared sequentially. He showed poor response to medical treatments and refused to receive advanced deep brain stimulation because of surgery risks. Five years after the first exhibition of symptoms and signs, the patient showed generalized and profound muscle twisting and contraction, including dysarthria and dysphagia. The patient presents sustained opitoshtonous-like posture and needs full assistance in his daily routines.

### Examination of Known Dystonia-Associated Mutations

Initial screening for known childhood-onset mutations on dystonia-associated genes *TOR1A, THAP1*, and ataxia-associated gene *FXN* identified a mutation of three-base pair deletion (c.907_909delGAG) in the patient’s *TOR1A* gene. This mutation has been previously reported to result in a pathogenic TOR1A (E303del) variant with an in-frame variant of Glu deletion (13). The patient’s *THAP1* and *FXN* genes are both normal alleles.

Additional screening in the core family members revealed that TOR1A (E303del) is also present in the genome of the patient’s father, but neither in his mother nor in sibling. This finding demonstrates that TOR1A (E303del) is originated from inheritance rather than de novo (**Figure 1A, 1B**). Notably, the patient and his core family members do not carry the protective dystonia-associated SNV (c.646G>C, p.D216H, rs1801968) on the TOR1A gene (14) (**Figure 1C, 1D**). This observation shows that the difference of symptom presentation between the patient and his father is not due to the protective role of TOR1A (D216H). Moreover, an expanded survey in other close family members further identified that the patient’s aunt and the first son of the aunt as asymptomatic carriers of TOR1A (E303del) (**Figure 1A, 1B**). These results collectively indicate that TOR1A (E303del) is insufficient to drive dystonia in this family.

**Figure 1.**
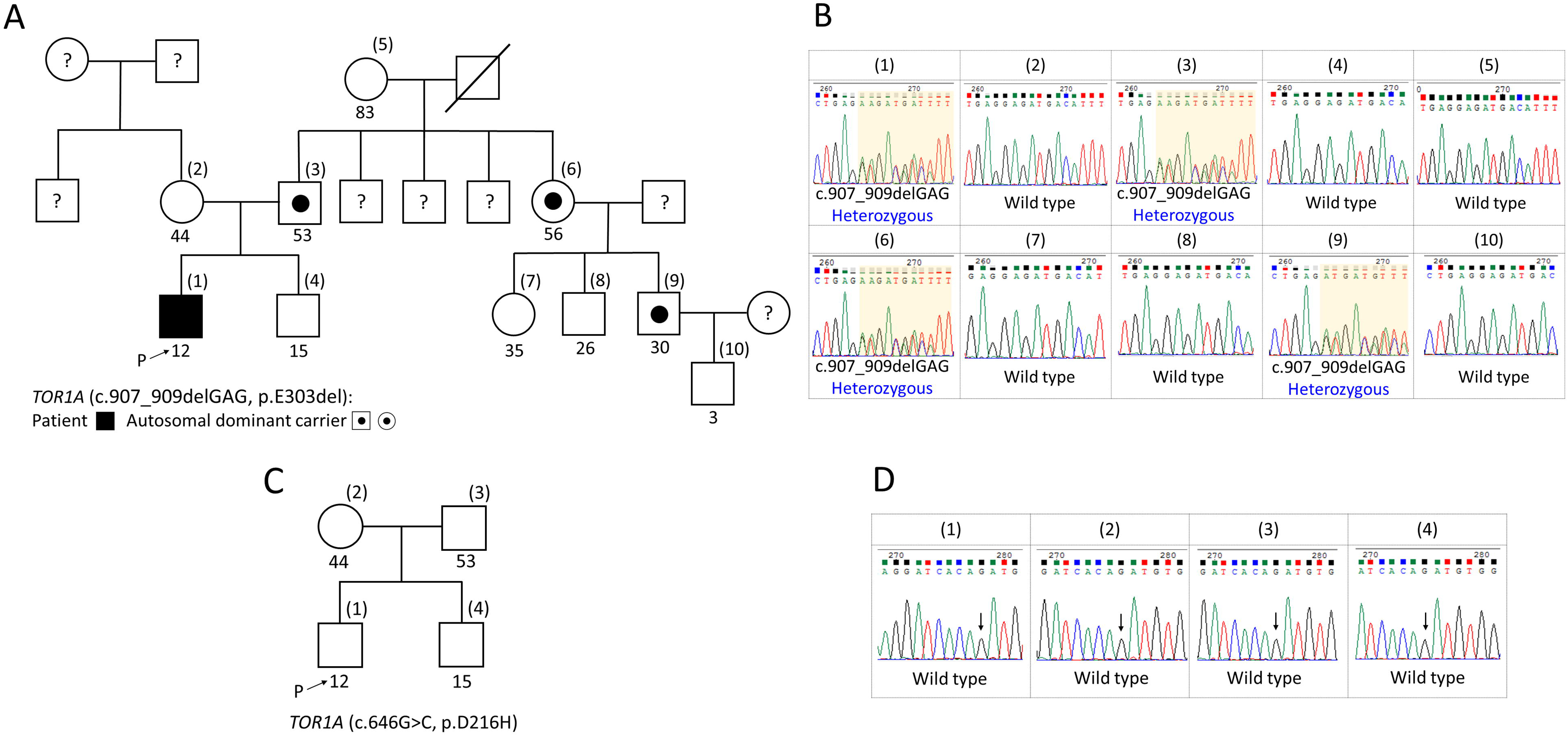
Family pedigree and Sanger sequencing data. Family pedigree **(A,C)** and Sanger sequencing data **(B,D)** of 10 family members **(A,B)** of *TOR1A* gene (c.907_909delGAG, p.E303del) and 4 family members (core family) **(C,D)** of *TOR1A* gene (c.646G>C, p.D216H). **(A,C)** The arrow point out the proband. The numbers within parentheses are the order of Sanger sequencing data and the numbers under the box/circle show the age (years old). The question marks within the box/circle indicate the unknown status because we don’t have the DNAs sample for study.

### Screening for Dystonia-Associated *TOR1A* Gene (c.907_909delGAG) Candidate Modifiers

#### All variants from *TOR1A* gene among the five WES data

WES was performed on the patient (subject 1) and three other TOR1A (E303del) carriers in the family to explore potential modifiers. Exomes of the patient’s mother (subject 2), who is neither a TOR1A (E303del) carrier nor symptomatic, was also examined as a contrast. While additional variants were found within the *TOR1A* gene of the mother, the father (subject 3), and the aunt (subject 6), including variants: C>T within promoter region (rs13300897), c.246G>A (p.A82A, rs2296793), and C>A on 3’-UTR (rs1182) (15,16), the patient and the first son of the aunt (subject 9) only have one genome variant *TOR1A* gene (c.907_909delGAG) in their *TOR1A* locus, which is pathogenic (**Table 1**).

**Table 1.**
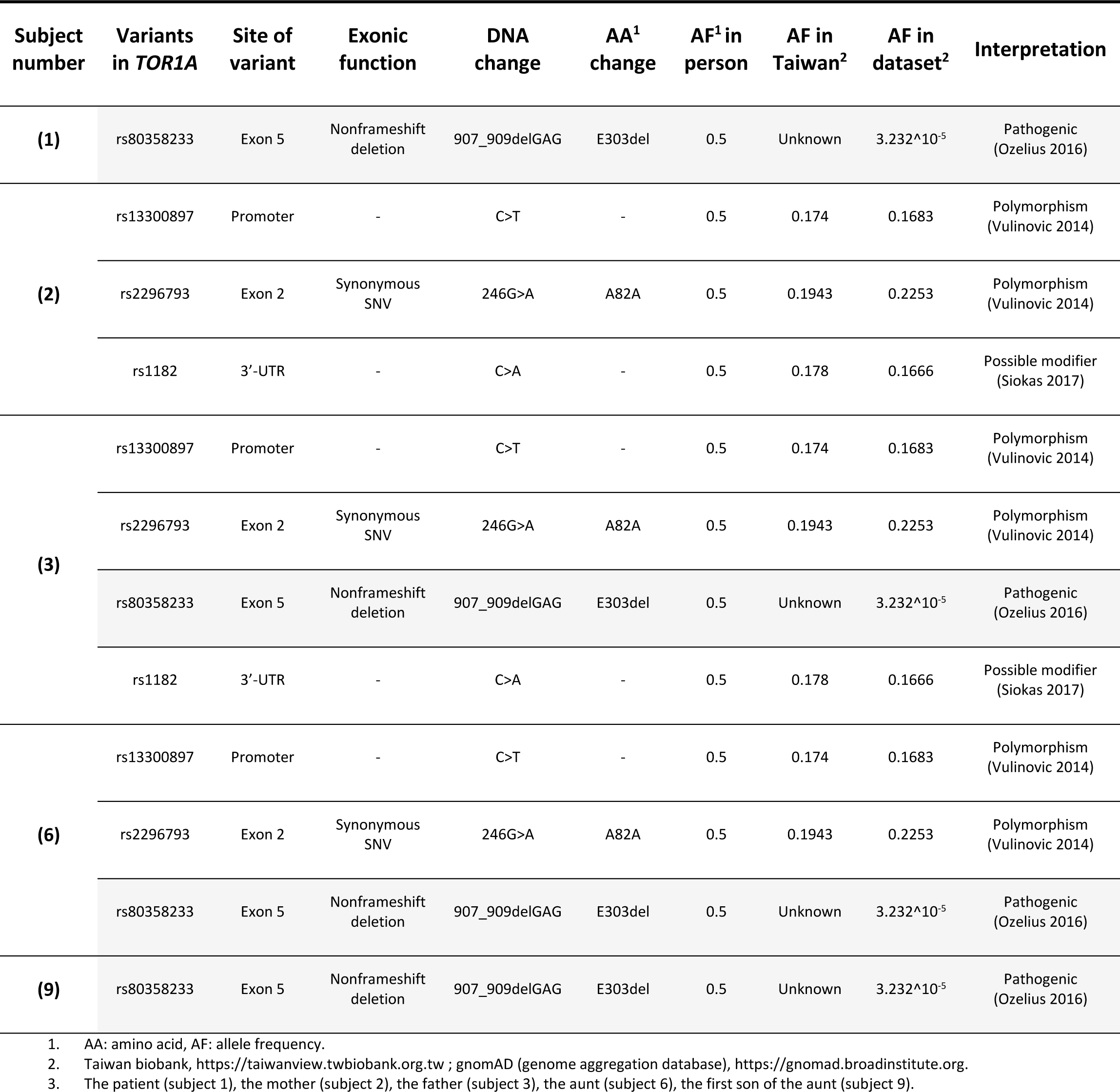
Genomic variants in the exons, promoter regions and 3’-UTR of the *TOR1A* gene between the patient and the other family members

#### Expand the search outside the *TOR1A* locus

We devised the following workflow to identify candidate genome variants from the five WES data systemically (**Figure 2**). After removing synonymous SNV, exonic genome variants are catalogued into three groups: (1) *de novo mutation* (*de novo*)--the allele frequency (AF) of the patient is 0.5 or 1 while all the other family members are 0; (2) *autosomal dominant inheritance* (AD)--the AF of the patient is 0.5 or 1 while the mother is 0.5 or 1 and the rest of the family members are 0; (3) *autosomal recessive inheritance* (AR)--the AF of the patient is 1 while the mother is 0.5 or 1, the father is 0.5, and both the aunt and the first son of the aunt are either 0 or 0.5. Subsequently, variants without clinical significance, predicted as tolerable and benign by SIFT and polyphen-2 respectively, and having sequencing depth less than 20 times were removed. With these filters, we identified 37 and 34 genome variants (**Supplementary Material_Table S1, S2**) in the AD group and AR group respectively, and none in the *de novo* group (**Figure 2**).

**Figure 2.**
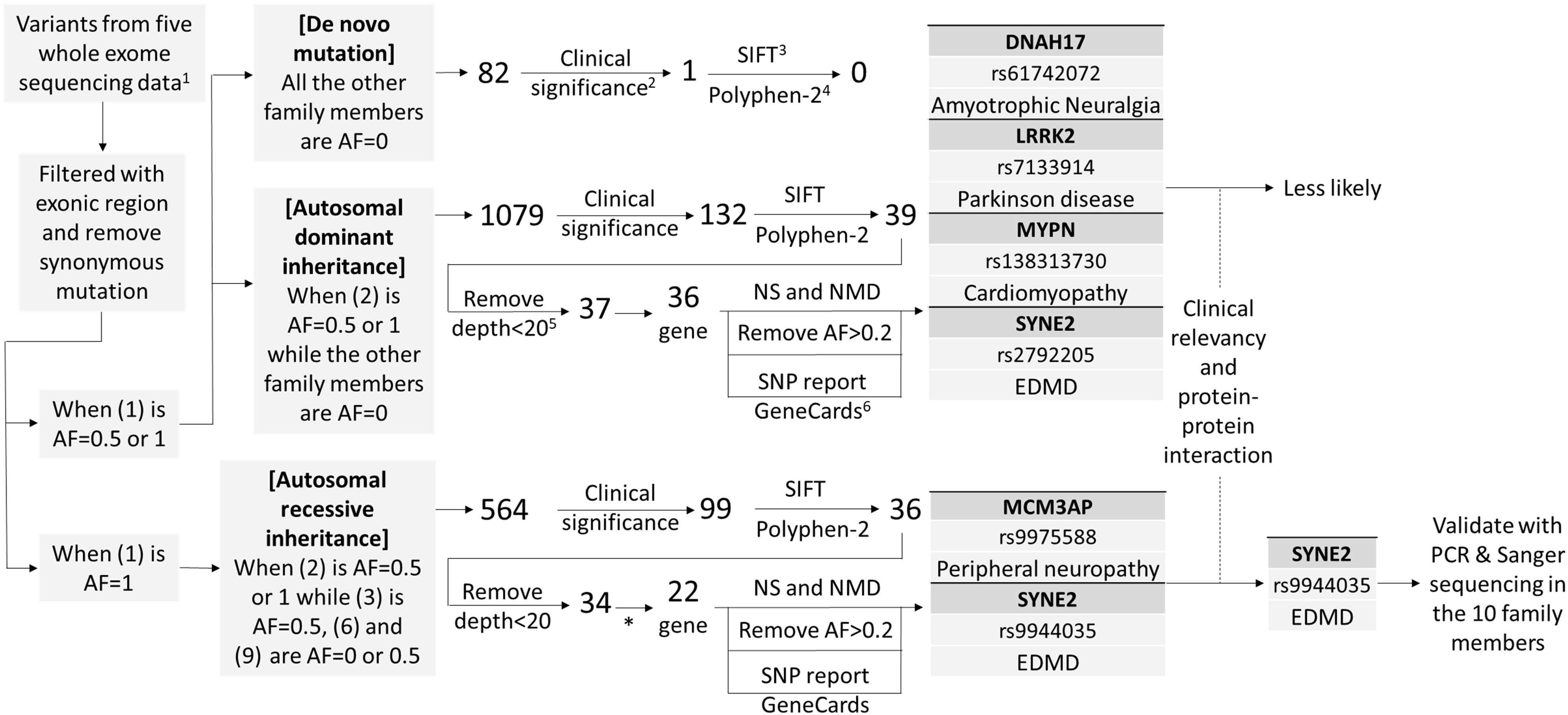
Workflow to find the candidate variants from either autosomal dominant or recessive inheritance. Data includes the patient (1), the mother (2), the father (3), the aunt (6), the first son of the aunt (9). Clinical significance: remove all the items without any annotation, SIFT: either deleterious or tolerated, Polyphen-2: benign, possibly damaging or damaging. Remove the variants with sequencing depth <20, Reference SNP report and human gene database-GeneCards**®** search to see the disease association and related publications. *: filtered out *TTN* gene (1 gene and 8 variants) at this step. AF (allele frequency), NS (neurologic disorders), NMD (neuromuscular diseases), EDMD (Emery-Dreifuss muscular dystrophy)

In the 34 variants from AR group, the *TTN* gene (containing 8 variants) was excluded for further consideration because of its repetitive sequence that may causes poor resolution on results from the next generation sequencing techniques (17). The remaining genes and variants were filtered with neurologic disorders and neuromuscular diseases firstly. Then, we removed the variants with AF higher than 0.2 which makes them tend to be tolerable polymorphism existing in the human population. Subsequently, we verified these variants by reference SNP reports and human gene database-GeneCards^®^ to review the clinical significance and publications. After critical review of clinical relevancy and extensive literature search for protein-protein interaction evidences, we firstly targeted on the variant, c.1721T>C (p.I574T, rs9944035) which resides in the *SYNE2* gene and we found it have been previously linked to Emery-Dreifuss muscular dystrophy (EDMD) in human disease (18).

#### Protein-protein interaction between SYNE2 and TOR1A

Notably, proteins encoded by the *SYNE2* gene (SYNE2) and *TOR1A* gene (TOR1A) physically interact with each other at the outer nuclear membrane (19). This finding suggests a potential genetic interaction between these genome variants in our patient. Thus, we validated the *SYNE2* gene (c.1721T>C) by Sanger sequencing in 10 family members and the result showed that the patient has homozygous mutation while the core family members and the grandmother have heterozygous alleles, which suggests its origin of inheritance. Moreover, homozygous *SYNE2* gene (c.1721T>C) was not found in other *TOR1A* gene (c.907_909delGAG) carriers within the family (**Figure 3**).

**Figure 3.**
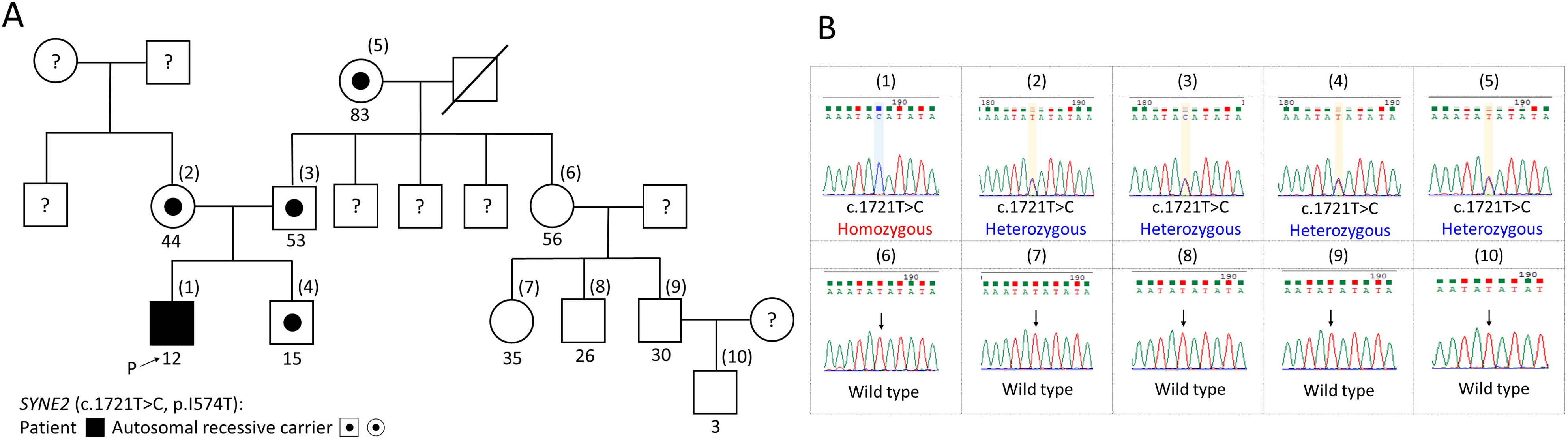
Family pedigree and Sanger sequencing data of 10 family members of *SYNE2* gene (c.1721T>C, p.I574T). Family pedigree **(A)** and Sanger sequencing data **(B)** of 10 family members of *SYNE2* gene (c.1721T>C). **(A)** The arrow points out the proband. The numbers within parentheses are the order of Sanger sequencing data and the numbers under the box/circle show the age (years old). The question marks within the box/circle indicate the unknown status because we don’t have the DNAs sample for study.

#### The potential role of SYNE2 on neurologic disorders

The *SYNE2* gene (c.1721T>C, p.I574T) is expected to result in an isoleucine-to-threonine change at the amino acid 574 located within the third spectrin repeat of the SYNE2. This isoleucine appears to be conserved among higher mammals, which provides an implication of the potential impact from having the variant (**Figure 4A, left**). Meanwhile, a great similarity of amino acid sequence between SYNE1 and SYNE2 implies *SYNE2* gene may have role in neurologic disorders as *SYNE1* gene does although no known annotation found in references yet (**Figure 4B**) (20).

**Figure 4.**
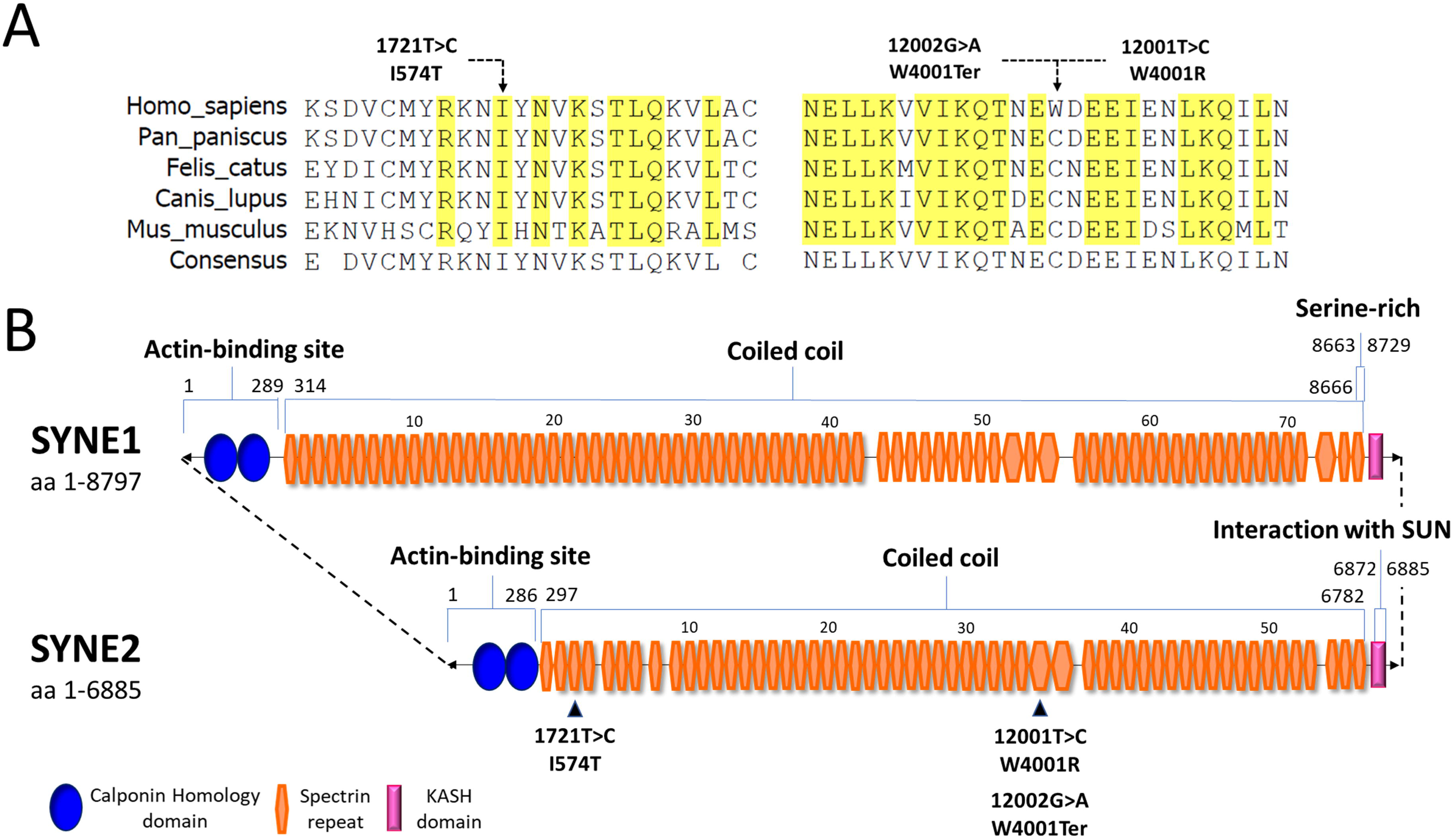
**(A)** Multiple alignment of SYNE2 protein (SYNE2 giant) on I574 and W4001 sites. Highlight areas show conserved amino acid sites through species. Sequences were aligned by using the VectorNTI tool. **(B)** Functional domain analysis between SYNE1 (NP_892006.3) and SYNE2 (NP_055995.4). Three major domains, calponin homology domain, spectrin repeat, and KASH domain, are labeled in blue, orange, and pink color.

#### Other hits from *SYNE2* gene may simultaneously enhance the disruption of protein-protein interaction

In addition to *SYNE2* gene (c.1721T>C), the patient has three more variants, c.12001T>C (p.W4001R, rs2792205), c.12002G>A (p.W4001Ter, rs2781377), and c.15556C>A (p.L5186M, rs10151658), found in the *SYNE2* coding region (**Table 2**). The incidence of *SYNE2* gene (c.15556C>A) is relatively high in Taiwan (0.6628) and this common prevalence renders it likely a polymorphism rather than a modifier. For *SYNE2* gene (c.12001T>C) and *SYNE2* gene (c.12002G>A), both mutations result in the amino acid change of residue W4001 into arginine and termination, SYNE2 (W4001R) and SYNE2 (W4001Ter), respectively. Although this amino acid, tryptophan, is a not conserved site among higher mammals (**Figure 4A, right**), we believe that SYNE2 (W4001R) and SYNE2 (W4001Ter) may quantitatively disrupt the physical interaction between TOR1A and SYNE2 at the outer nuclear membrane as well as SYNE2 (I574T) does.

**Table 2.** Genomic variants in the exons of the *SYNE2* gene between the patient and the other family members

| Subject number | Variants in SYNE2 | Exonic function | DNA change | AA change | AF in person | AF in Taiwan | AF in dataset | Clinical significance<sup>1</sup> | SIFT | Polyphen2 |
| --- | --- | --- | --- | --- | --- | --- | --- | --- | --- | --- |
| (1) | rs10151658 | Nonsynonymous SNV | 15556C>A | L5186M | 1 | 0.6628 | 0.5669 | Benign | Tolerable | Benign |
|  | rs9944035 | Nonsynonymous SNV | 1721T>C | I574T | 1 | 0.1171 | 0.0867 | Benign | Deleterious | Benign |
|  | rs2792205 | Nonsynonymous SNV | 12001T>C | W4001R | 0.5 | 0.1101 | 0.0815 | Benign | Deleterious | Benign |
|  | rs2781377 | Stop-gain | 12002G>A | W4001Ter | 0.5 | 0.1049 | 0.0811 | Benign | Unknown | Unknown |
| (2) | rs10151658 | Nonsynonymous SNV | 15556C>A | L5186M | 0.5 | 0.6628 | 0.5669 | Benign | Tolerable | Benign |
|  | rs9944035 | Nonsynonymous SNV | 1721T>C | I574T | 0.5 | 0.1171 | 0.0867 | Benign | Deleterious | Benign |
|  | rs2792205 | Nonsynonymous SNV | 12001T>C | W4001R | 1 | 0.1101 | 0.0815 | Benign | Deleterious | Benign |
|  | rs2781377 | Stop-gain | 12002G>A | W4001Ter | 1 | 0.1049 | 0.0811 | Benign | Unknown | Unknown |
| (3) | rs10151658 | Nonsynonymous SNV | 15556C>A | L5186M | 0.5 | 0.6628 | 0.5669 | Benign | Tolerable | Benign |
|  | rs9944035 | Nonsynonymous SNV | 1721T>C | I574T | 0.5 | 0.1171 | 0.0867 | Benign | Deleterious | Benign |
|  | rs37378453<br>3 | Nonsynonymous SNV | 9034T>A | W3012R | 0.5 | Unknown | 3.229<sup>^</sup>10<sup>-5</sup> | Unknown | Deleterious | Damaging |
| (6) | rs13868905<br>3 | Nonsynonymous SNV | 9347A>G | K3116R | 0.5 | 0.0092 | 0.0024 | Benign | Tolerable | Benign |
| (9) | rs10151658 | Nonsynonymous SNV | 15556C>A | L5186M | 0.5 | 0.6628 | 0.5669 | Benign | Tolerable | Benign |

1. Clinical significance from refence SNP report

## Discussion

While homozygous mutations in DYT1 dystonia lead to profound morbid, even lethal, in both human and mouse model (21,22), the fact that most patients carrying heterozygous mutations with variable disease penetrance suggests potential involvement of genetic modifiers. Numerous attempts have been made to understand the low penetrance rate of this disease in a heterozygous form. For example, it is known that interaction of TOR1A with its major binding partners, TOR1AIP1 and 2 or HSPA8, are impaired by the dystonia-associated GAG Deletion (8,9). However, the patient in our study does not have the variants in his *TOR1AIP2* and *HSPA8* genes. Despite he does carry two variants, c.437T>C (p.M146T, rs1281378) and c.827C>G (p.P276R, rs609521), in the *TOR1AIP1* gene, the high AF in Taiwan-based population (both greater than 0.7) suggests that they are less likely as culprits in our case.

Since DYT1 dystonia is categorized as a CNS disease and best conceptualized as a motor circuit disorder (23), we focused primarily on neurologic disorders and neuromuscular diseases to find out the potential modifier variants. After advanced filtering, we propose that SYNE2 (I574T) could be the candidate variant initially. *SYNE2* gene is associated with EDMD, which is a condition that primarily affects skeletal muscles. The earliest features of EDMD are joint contractures, which restrict the movement of certain joints and usually become noticeable in early childhood (24). The most striking and intriguing finding is TOR1A and SYNE2 physically interact with each other and exert vital biologic functions (19). The TOR1A encoded by *TOR1A* gene binds the KASH domain of SYNE (nesprin) and participates in linkage between nuclear envelope and cytoskeleton (**Figure 5**).

**Figure 5.**
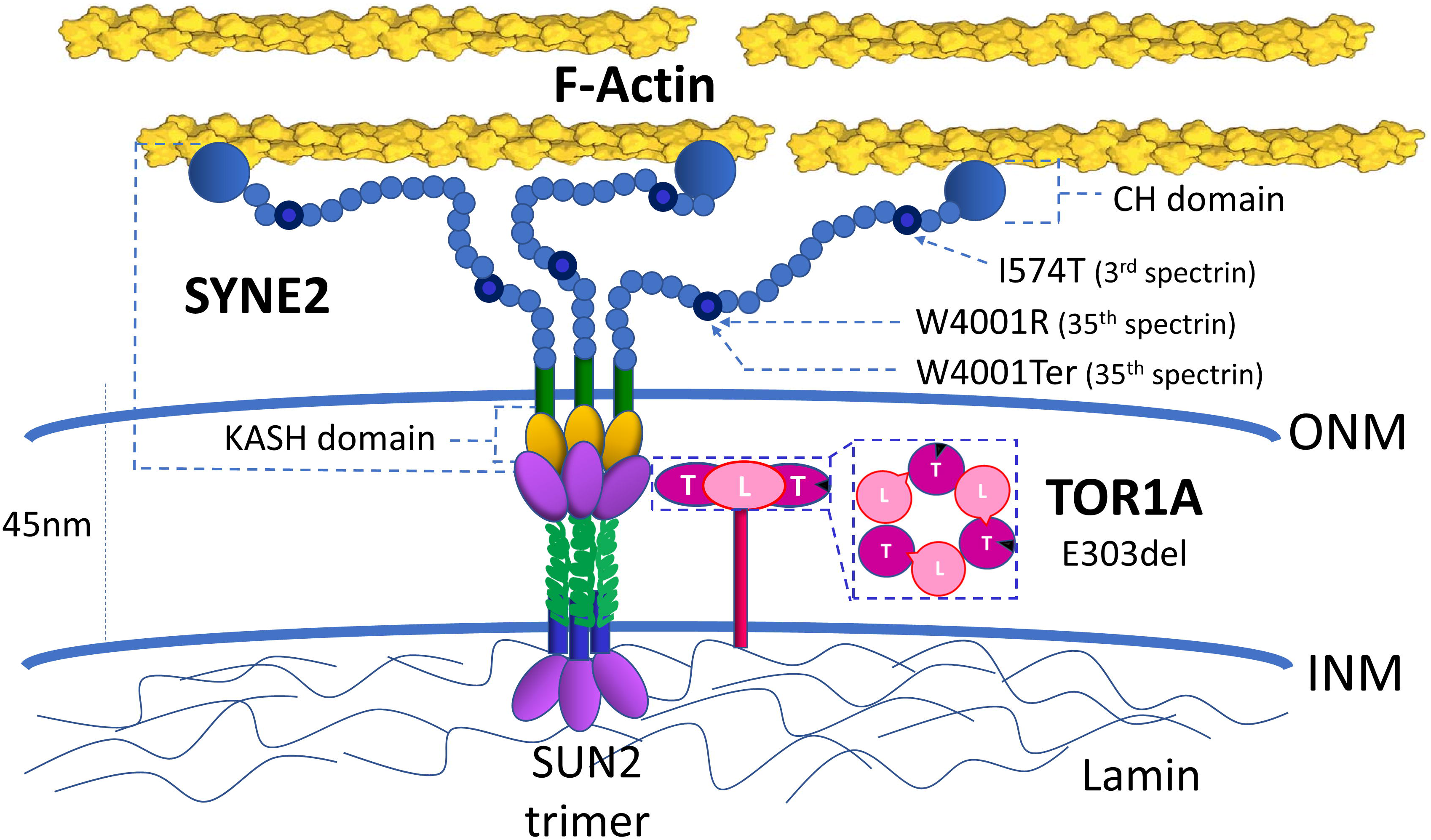
TOR1A-LAP1 (or LULL1) heterohexamer regulates the assembly and function of LINC complex. The location of the defects at TOR1A (E303del) and at SYNE2 (3^rd^ spectrin, I574T; 35^th^ spectrin: W4001R and W4001Ter). LINC complex (the linker of nucleoskeleton and cytoskeleton, consisting of KASH domain and SUN proteins), T (TOR1A), L (LAP1 or LULL1), CH domain (Calponin Homology domain), KASH domain (Klarsicht, ANC-1, and Syne homology domain), SUN2 (SUN (Sad1, UNC-84) domain-containing protein 2), ONM (outer nuclear membrane), INM (inner nuclear membrane).

SYNE are a family of proteins that are found primarily in the outer nuclear membrane and are part of the LINC (linker of nucleoskeleton and cytoskeleton) complex (25). The SYNE2 (I574T) variant is at the third spectrin repeat in SYNE2 and spectrin is an important mechanoresponsive protein shaping fusogenic synapse architecture during myoblast fusion (26). Not only the SYNE2 (I574T) variant is locally close to the Calponin Homology domain of SYNE2 where is a pivotal area for actin binding (**Figure 5**), but also the impact on a highly conserved amino acid in the third spectrin repeat suggests a functional significance of this variant. Furthermore, more hits on SYNE2 (W4001R) and SYNE2 (W4001Ter) respectively found in this case may seriously disrupt the physical interaction between TOR1A and SYNE2 at the same time. Thus, we believe these three variants found within the *SYNE2* gene may play the vital role on accelerating disease onset.

Saunders and Luxton elaborated how defects in LINC complex regulation by TOR1A may contribute to the pathogenesis of DYT1 dystonia although the precise regulatory mechanism remains unclear (27). Intriguingly, the loss of the glutamic acid residue in the C-terminal of one or more subunits of TOR1A might act to disrupt the interaction with a partner protein (LAP1 or LULL1) or the closure of the ring (heterohexamer) (**Figure 5**) (28), but this is not sufficient to cause DYT1 disease onset by dominant effects of TOR1A (E303del). Another hits on the 3^rd^ and 35^th^ spectrin of SYNE2 may seriously put the LINC complex into profound dysfunction and clinically accelerate the disease onset. In view of LINC complex-dependent molecular bridge for physically coupling the nucleus to the cytoskeleton, it may come without surprise that mutations in the genes encoding SYNE and SUN proteins (**Figure 5**) are associated with an ever-expanding list of human diseases, including ataxia and muscular dystrophy (27). Furthermore, we noticed the amino acids sequence similarity between SYNE1 and SYNE2 and they both belong to nesprin family. Since the dysfunction of *SYNE1* gene is associated with spinocerebellar ataxia, we believe that *SYNE2* gene may have role on neurologic disorders (19).

The other variants that are associated with neurologic disorders and neuromuscular diseases (reviewed by human gene database-GeneCard^®^), like *DNAH17* gene (c.11857C>T, p.H3953Y, rs61742072), *LRRK2* gene (c.4193G>A, p.R1398H, rs7133914), *MYPN* gene (c.3481C>A, p.L1161I, rs138313730), and *MCM3AP* gene (c.305C>T, p.S102L, rs9975588) also emerged in our candidate list (**Figure 2**). However, these variants less likely serve as the TOR1A (E303del) modifiers due to lack of protein-protein interaction evidence after meticulous and broad literature search and review. Regarding the limitation of this article, the basic issue is that we need more patients with TOR1A (E303del) to testify our expectation whether variants in SYNE2 play a role in DYT1 penetrance. We also need functional assessment in the future experimentation in mammalian species to clarify their roles and contributions in the DYT1 dystonia.

## Conclusion

In summary, we propose that SYNE2 variants maybe the potential modifier SNVs which could drop the threshold of disease onset of DYT1 dystonia and facilitates the clinical symptoms and signs of dystonia. We believe that this study provided a clue to unravel the candidate SNVs and try to find the potential modifier variants from this family. Our findings not only echo the previous research highlighting the KASH-SUN interaction and LINC complex regulation by TOR1A, but provide knowledge for further understanding the disease origin of the DYT1 dystonia as well. We will recommend the physicians to test these variants once the *TOR1A* gene (c.907_909delGAG) patient show normal alleles within other *TOR1A* locus and other major binding proteins, such as *SYNE2* gene in this study.

## Ethics Statement

This study was approved by the Ethics Committee of Tri-Service General Hospital, National Defense Medical Center in Taiwan, which was in accordance with the ethical standards of the institutional and/or national research committee and with the 1964 Helsinki declaration and its later amendments or comparable ethical standards. And written informed consent was obtained from all subjects. We also obtained written and informed consent from the patients who gave specific permission to publish the data.

## Authors Contributions

FCL and CFH: conceptualization; FCL and CFH: methodology; CSH: software; SW: validation; FCL and CSH: formal analysis; FCL and SW: investigation; SW and JSH: resources; FCL and CSH: data curation; FCL, SW, and SMH: writing-original draft; CFH: writing-review and editing; SMH and CFH: supervision; CFH: project administration; CFH: funding acquisition.

## Acknowledgements

We would like to thank the patient and his family members who provided the DNAs and clinical information necessary for this research study.

## References

1. Bressman SB, de Leon D, Kramer PL, Ozelius LJ, Brin MF, Greene PE, et al. Dystonia in Ashkenazi Jews: clinical characterization of a founder mutation. Ann Neurol. (1994) 36:771–777. doi.org/10.1002/ana.410360514

2. Fernández-Alvarez E, Nardocci N. Update on pediatric dystonias: etiology, epidemiology, and management. Degener Neurol Neuromuscul Dis. (2012) 11:29–41. doi: 10.2147/DNND.S16082.

3. Ozelius LJ, Bressman SB. Genetic and clinical features of primary torsion dystonia. Neurobiol Dis. (2011) 42:127–35. doi.org/10.1016/j.nbd.2010.12.012

4. Jahanshahi M, Torkamani M. The Cognitive Features of Idiopathic and DYT1 Dystonia. Mov Disord. (2017) 32:1348–1355. doi.org/10.1002/mds.27048

5. Ozelius LJ, Hewett JW, Page CE, Bressman SB, Kramer PL, Shalish C, et al. The early-onset torsion dystonia gene (DYT1) encodes an ATP-binding protein. Nat Genet. (1997) 17:40–8. doi.org/10.1038/ng0997-40

6. Charlesworth G, Bhatia KP, Wood NW. The genetics of dystonia: new twists in an old tale. Brain. (2013) 136:2017–2037. doi.org/10.1093/brain/awt138

7. Walter M, Bonin M, Pullman RS, Valente EM, Loi M, Gambarin M, et al. Expression profiling in peripheral blood reveals signature for penetrance in DYT1 dystonia. Neurobiol Dis. (2010) 38(2):192–200. doi.org/10.1016/j.nbd.2009.12.019.

8. Leung JC, Klein C, Friedman J, Vieregge P, Jacobs H, Doheny D, et al. Novel mutation in the TOR1A (DYT1) gene in atypical early onset dystonia and polymorphisms in dystonia and early onset parkinsonism. Neurogenetics. (2001) 3(3): 133–43. doi.org/10.1007/s100480100111

9. Calakos N, Patel VD, Gottron M, Wang G, Tran-Viet KN, Brewington D, et al. Functional evidence implicating a novel TOR1A mutation in idiopathic, late-onset focal dystonia. J Med Genet. (2010) 47(9):646–50. doi.org/10.1136/jmg.2009.072082.

10. Naismith TV, Dalal S, Hanson PI. Interaction of torsinA with its major binding partners is impaired by the dystonia associated DeltaGAG deletion. J Biol Chem. (2009) 284(41):27866–74. doi.org/10.1074/jbc.M109.020164

11. Siokas V, Aloizou AM, Tsouris Z, Michalopoulou A, Mentis AFA, Dardiotis E. Risk Factor Genes in Patients with Dystonia: A Comprehensive Review. Tremor Other Hyperkinet Mov. (2019) 8:559. doi.org/10.7916/D8H438GS

12. DePristo MA, Banks E, Poplin R, Garimella KV, Maguire JR, Hartl C, et al. A framework for variation discovery and genotyping using next-generation DNA sequencing data. Nat Genet. (2011) 43(5):491–8. doi.org/10.1038/ng.806

13. Warner TT, Jarman P. The molecular genetics of the dystonias. J Neurol Neurosurg Psychiatry. (1998) 64:427–9. doi.org/10.1136/jnnp.64.4.427

14. Kamm C, Fischer H, Garavaglia B, Kullmann S, Sharma M, Schrader C, et al. Susceptibility to DYT1 dystonia in European patients is modified by the D216H polymorphism. Neurology. (2008) 70(23):2261–2. doi.org/10.1212/01.wnl.0000313838.05734.8a

15. Siokas, V.; Dardiotis, E.; Tsironi, E.E.; Tsivgoulis, G.; Rikos, D.; Sokratous, M.; et al. The Role of TOR1A Polymorphisms in Dystonia: A Systematic Review and Meta-Analysis. PLoS One. 2017, 12(1), e0169934. doi.org/10.1371/journal.pone.0169934.

16. Vulinovic F, Lohmann K, Rakovic A, Capetian P, Alvarez-Fischer D, Schmidt A, et al. Unraveling cellular phenotypes of novel TorsinA/TOR1A mutations. Hum Mutat. (2014) 35(9):1114–22. doi.org/10.1002/humu.22604

17. Hackman P, Evila A, Udd B. G.P.17: TTN a challenge for next generation sequencing. Neuromuscular Disorders. (2014) 24(9–10):799. doi.org/10.1016/B978-0-444-59565-2.00007-1

18. Zhang Q, Bethmann C, Worth NF, Davies JD, Wasner C, Feuer A, et al. Nesprin-1 and −2 are involved in the pathogenesis of Emery-Dreifuss muscular dystrophy and are critical for nuclear envelope integrity. Hum Mol Genet. (2007) 16:2816–2833. doi.org/10.1016/B978-0-444-59565-2.00007-1

19. Nery FC, Zeng J, Niland BP, Hewett J, Farley J, Irimia D, et al. TorsinA binds the KASH domain of nesprins and participates in linkage between nuclear envelope and cytoskeleton. J Cell Sci. (2008) 121(Pt 20):3476–86. doi.org/10.1242/jcs.029454

20. Peng Y, Ye W, Chen Z, Peng H, Wang P, Hou X, et al. Identifying SYNE1 Ataxia With Novel Mutations in a Chinese Population. Front Neurol. (2018) 9:1111. doi.org/10.3389/fneur.2018.01111

21. Goodchild RE, Kim CE, Dauer WT. Loss of the dystonia-associated protein torsinA selectively disrupts the neuronal nuclear envelope. Neuron (2005) 48:923–932. doi.org/10.1016/j.neuron.2005.11.010

22. Kariminejad A, Dahl-Halvarsson M, Ravenscroft G, Afroozan F, Keshavarz E, Goullée H, et al. TOR1A variants cause a severe arthrogryposis with developmental delay, strabismus and tremor. Brain. (2017) 140(11):2851–2859. doi.org/10.1093/brain/awx230

23. Tanabe LM, Kim CE, Alagem N, Dauer WT. Primary dystonia: molecules and mechanisms. Nat Rev Neurol. (2009) 5(11):598–609. doi.org/10.1038/nrneurol.2009.160

24. Bonne G, Leturcq F, Ben Yaou R. Emery-Dreifuss Muscular Dystrophy. GeneReviews®. 2015. Seattle (WA): University of Washington, Seattle, 1993-2019.

25. Rajgor D, Shanahan CM. Nesprins: from the nuclear envelope and beyond. Expert Rev Mol Med. (2013) 15, e5. doi.org/10.1017/erm.2013.6

26. Duan R, Kim JH, Shilagardi K, Schiffhauer ES, Lee DM, Son S, et al. Spectrin is a mechanoresponsive protein shaping fusogenic synapse architecture during myoblast Fusion. Nat Cell Biol. (2018) 20(6):688–698. doi.org/10.1038/s41556-018-0106-3

27. Saunders CA, Luxton GW. LINCing Defective Nuclear-Cytoskeletal Coupling and DYT1 Dystonia. Cell Mol Bioeng. (2016) 9(2):207–216. doi.org/10.1007/s12195-016-0432-0

28. Breakefield XO, Kamm C, Hanson PI. TorsinA: movement at many levels. Neuron. (2001) 31(1):9–12. doi.org/10.1016/S0896-6273(01)00350-6

