## Supplementary_Material Tables 1-2 for "Association of SYNE2 variants in accelerating the progress of DYT1 early-onset isolated dystonia"

### Supplementary Tables 1-2

**Table S1. Genomic variants in the exons and nonsynonymous change in autosomal dominant inheritance model (37 variants/36 genes)**

| Hits number | Subject links | Variants | Gene symbol | DNA change | AA change | AF in Taiwan | AF in dataset | Clinical significance | SIFT | Polyphen2 |
|---|---|---|---|---|---|---|---|---|---|---|
| 2 | 1,2 | rs188159332 | *ANK3* | G4907A | R1636K | 0.0069 | 0.0003 | Not specified | T | D |
| | | rs3858340 | *BAG3* | C1220T | P407L | 0.2503 | 0.1263 | Benign | D | B |
| | | rs116952709 | *BRIP1* | G430A | A144T | 0.0228 | 0.0013 | Benign | T | D |
| | | rs3753058 | *CD3G* | G391T | V131F | 0.4859 | 0.1723 | Benign | D | P |
| | | rs3735972 | *CNGB3* | A2264G | E755G | 0.0757 | 0.0887 | Benign | D | B |
| | | rs142433421 | *COG5* | T1856C | I619T | 0.0238 | 0.0016 | Likely benign | D | B |
| | | rs61742072 | *DNAH17* | C11857T | H3953Y | 0.1040 | 0.1201 | Benign | D | B |
| | | rs1990236 | *DNAH9* | G2058A | M686I | 0.1972 | 0.1757 | Benign | D | B |
| | | rs115095929 | *DRC1* | A2147C | Q716P | 0.0482 | 0.0239 | Benign | D | P |
| | | rs146567337 | *EDAR* | A1138C | S380R | 0.0241 | 0.0009 | Benign | D | D |
| | | rs11570255 | *EDN3* | G49A | A17T | 0.0149 | 0.0015 | Benign | T | P |
| | | rs11569017 | *EGF* | A2225T | D742V | 0.1881 | 0.0564 | Likely benign | D | B |
| | | rs2302427 | *EZH2* | G436C | D146H | 0.1778 | 0.0635 | Benign | D | P |
| | | rs6183 | *GHR* | C1417A | P473T | 0.0343 | 0.0013 | Likely benign | D | D |
| | | rs696217 | *GHRL* | C178A | L60M | 0.1791 | 0.0721 | Pathogenic | T | D |
| | | rs2274084 | *GJB2* | G79A | V27I | 0.3072 | 0.0222 | Benign | T | D |
| | | rs145762716 | *GLMN* | C761G | P254R | 0.0152 | 0.0010 | Likely benign | T | P |

| | | | | | | | | | | |
|---|---|---|---|---|---|---|---|---|---|---|
| | | rs2296434 | *HPS1* | C1112G | P371R | 0.1827 | 0.1089 | Benign | T | D |
| | | rs1050349 | *LAMA4* | C3356G | P1119R | 0.3734 | 0.2140 | Benign | T | D |
| | | rs1048661 | *LOXL1* | G422T | R141L | 0.4472 | 0.2890 | Not specified | D | B |
| | | rs2306029 | *LRP4* | A4660G | S1554G | 0.2422 | 0.4283 | Benign | D | B |
| | | rs9297853 | *LRRC6* | T1151C | I384T | 0.0536 | 0.1485 | Benign | D | P |
| | | rs7133914 | *LRRK2* | G4193A | R1398H | 0.0881 | 0.0900 | Not specified | T | P |
| | | rs3211097 | *MFF* | A19T | S7C | 0.2363 | 0.2736 | Benign | D | B |
| | | rs3817552 | *MYBPC1* | C1365G | H455Q | 0.2942 | 0.1520 | Benign | D | D |
| | | rs138313730 | *MYPN* | C3481A | L1161I | 0.0056 | 0.0018 | Likely benign | D | D |
| | | rs61755995 | *NIN* | C2698T | R900C | 0.1185 | 0.0080 | Likely benign | D | P |
| | | rs56128139 | *NME8* | T1478C | I493T | 0.1191 | 0.2575 | Benign | D | B |
| | | rs114615449 | *NPHS1* | G1802C | G601A | 0.0176 | 0.0012 | Not specified | D | D |
| | | rs28939695 | | G1339A | E447K | 0.0281 | 0.0015 | Pathogenic | T | D |
| | | rs121918027 | *PLG* | G1858A | A620T | 0.0152 | 0.0009 | Pathogenic | T | D |
| | | rs357564 | *PTCH1* | C3944T | P1315L | 0.5153 | 0.3355 | Benign | D | P |
| | | rs259290 | *RGS9BP* | G286T | A96S | 0.2730 | 0.5412 | Benign | T | P |
| | | rs1804495 | *SERPINA7* | G909T | L303F | 0.2545 | 0.1127 | Pathogenic | D | P |
| | | rs2792205 | *SYNE2* | T12001C | W4001R | 0.1101 | 0.0815 | Benign | D | B |
| | | rs3744438 | *TBX4* | C941T | A314V | 0.1074 | 0.1904 | Benign | D | B |
| | | rs12498609 | *TET2* | C86G | P29R | 0.1985 | 0.0411 | Not specified | D | P |

1. SIFT: either deleterious (D) or tolerated (T).
2. Polyphen-2: benign (B), possibly damaging (P) or damaging (D).
3. Subject 1: the patient, subject 2: the mother.

| **Table S2. Genomic variants in the exons and nonsynonymous change in autosomal recessive inheritance model (26 variants/22 genes after minus *TTN* gene with 8 variants)** | | | | | | | | | | |
|---|---|---|---|---|---|---|---|---|---|---|
| **Hits number** | **Subject links** | **Variants** | **Gene symbol** | **DNA change** | **AA change** | **AF in Taiwan** | **AF in dataset** | **Clinical significance** | **SIFT** | **Polyphen2** |
| 3 | 1,2,3 | rs17853192 | *CPA6* | C518G | S173C | 0.2226 | 0.0721 | Benign | D | P |
| | | rs1799930 | *NAT2* | G590A | R197Q | 0.2599 | 0.2702 | Drug response | T | D |
| | | rs4715 | *SFTPC* | C254A | T85N | 0.2912 | 0.2167 | Benign | T | P |
| | | rs344141 | *SHROOM3* | C1405G | P469A | 0.2388 | 0.5261 | Benign | D | D |
| | | rs9944035 | *SYNE2* | T1721C | I574T | 0.1171 | 0.0867 | Benign | D | B |
| 4 | 1,2,3, 6 | rs12607385 | *CFAP53* | C691T | R231C | 0.4462 | 0.0402 | Benign | D | B |
| | | rs4935502 | *PCDH15* | A1193C | D398A | 0.1549 | 0.1769 | Benign | D | D |
| | | rs2228570 | *VDR* | T152C | M51T | 0.4658 | 0.6483 | Benign | D | B |
| | 1,2,3, 9 | rs885479 | *MC1R* | G488A | R163Q | 0.3869 | 0.1022 | Benign | D | B |
| | | rs1800866 | *SIGMAR1* | A5C | Q2P | 0.3251 | 0.1684 | Not specified | T | P |
| 5 | 1,2,3, 6,9 | rs4961 | *ADD1* | G1378T | G460W | 0.4531 | 0.1700 | Drug response | D | D |
| | | rs4916685 | *ADGRV1* | C5960T | P1987L | 0.3767 | 0.3106 | Not specified | T | D |
| | | rs10037067 | | A6695G | Y2232C | 0.3779 | 0.3204 | Not specified | D | P |
| | | rs2366926 | | A7034G | N2345S | 0.3758 | 0.2989 | Not specified | T | D |
| | | rs3784678 | *C15orf41* | C106G | L36V | 0.3580 | 0.4493 | Benign | T | P |
| | | rs2285944 | *DNAH11* | A101T | E34V | 0.2732 | 0.4433 | Benign | D | B |
| | | rs16895517 | *EYS* | C5617G | L1873V | 0.1780 | 0.0937 | Benign | T | P |
| | | rs62415827 | | C4543T | R1515W | 0.1800 | 0.0951 | Benign | D | P |
| | | rs448012 | *FLT4* | C2670G | H890Q | 0.4656 | 0.6216 | Benign | D | B |
| | | rs2229519 | *GBE1* | A568G | R190G | 0.4587 | 0.3263 | Benign | D | B |
| | | rs14024 | *KRT1* | A1898G | K633R | 0.3989 | 0.2661 | Benign | D | B |

| | | | | | | | | | | |
|---|---|---|---|---|---|---|---|---|---|---|
| | | rs9975588 | *MCM3AP* | C305T | S102L | 0.1287 | 0.3071 | Benign | D | B |
| | | rs2287780 | *MTRR* | C1243T | R415C | 0.1819 | 0.0525 | Benign | D | D |
| | | rs16879334 | | C1349G | P450R | 0.1814 | 0.0480 | Benign | D | D |
| | | rs1800566 | *NQO1* | C343T | P115S | 0.4706 | 0.2066 | Drug response | T | D |
| | | rs4148323 | *UGT1A1* | G211A | G71R | 0.1589 | 0.0149 | Drug response | D | B |

1. SIFT: either deleterious (D) or tolerated (T).

2. Polyphen-2: benign (B), possibly damaging (P) or damaging (D).

3. Subject 1: the patient, subject 2: the mother, subject 3: the father, subject 6: the aunt, subject 9: the first son of the anut.
